# Niacin protects against vascular calcification via enhanced SIRT1 and SIRT6 signaling and promoted VSMC autophagy

**DOI:** 10.1101/2025.01.07.631823

**Authors:** Chao-hua Kong, Li-da Wu, Yue Sun, Xiao-min Jiang, Yi Shi, Feng Wang, Dong-chen Wang, Yue Gu, Wen-ying Zhou, Jin-que Luo, Yue-lin Chao, Shao-liang Chen

## Abstract

**Background:** Vascular calcification (VC) is a common pathologic state that often accompanies calcium-phosphorus metabolism disorder and chronic kidney diseases (CKDs) in aging individuals. Vascular smooth muscle cell (VSMC) has been widely acknowledged as one of the main cell types that involved in this process. Niacin, a lipid-lowering reagent, has been demonstrated to be beneficial in atherosclerotic disease. The correlation and mechanism between niacin and vascular calcification have not been reported so far.

**Methods and results:** RCS analysis based on large clinical databases indicated that diet that riches in niacin can protect against abdominal artery calcification (AAC). Our data showed that niacin treatment remarkably reduced VSMC osteogenic differentiation and senescence-associated markers under osteogenic stimulation. Moreover, niacin treatment alleviated CKD and vitamin D_3_-induced vascular calcification in C57BL/6J mice. Mechanistically, we for the first time demonstrated that niacin supplementation inhibited vascular calcification via maintaining both Sirtuin 1 (SIRT1) and Sirtuin 6 (SIRT6) levels. Moreover, we verified that niacin activated SIRT1 and SIRT6 promoted VSMC autophagy flux.

**Conclusions:** These findings may help to develop novel therapeutic strategies in the treatment and prevention of vascular calcification.

## Introduction

Vascular calcification (VC) is characterized by the deposition of calcium-phosphate complexes in the tunica media and is considered as a degradative process associated with natural aging process, genetic diseases and certain pathological process such as diabetes, CKDs and hypertension^1-3^. Indeed, the vasculature system is the second most calcified tissue after the skeleton and a large number of studies have suggested that VC is a gene-related transdifferentiation process resembling bone mineralization^4^. Accumulating evidences have convinced that vascular smooth muscle cell (VSMC) osteoblast-like dedifferentiation plays the most essential pathological role in VC. Mechanisms associated with VC including inflammation, cellular autophagy, endoplasmic reticulum stress, cellular senescence and aberrant calcium-phosphate metabolism^5-8^. Targeting the osteogenic transition of VSMC becomes an important and promising strategy in the prevention and treatment of vascular calcification.

Vascular calcification features vascular stiffness, is a process usually accompanies cellular senescence. Of note, Vascular aging is the fundamental pathophysiological factor for various cardiovascular diseases including atherosclerosis, vascular calcification and aneurysm^9^. Senescent VSMC exhibits a long and narrow look and reduced contractile ability with accelerated trend of osteogenic transition upon pathological stimulation^10^. The Sirtuin family comprises 7 members and is known to play important role in regulating cellular senescence^11^. Interestingly, previous studies have indicated the Sirtuin proteins are involved in vascular calcification especially Sirtuin 1 and Sirtuin 6^12,13^. Both Sirtuin 1 and Sirtuin 6 are nicotinamide adenine dinucleotide-dependent histone deacetylases that involved in regulating cell proliferation, senescence, differentiation and apoptosis and are reduced in calcified vascular^14,15^. Accumulating evidences have indicated that Sirtuin 1 and Sirtuin 6 play critical role in age-associated pathological alterations including cardiovascular diseases, metabolic disease and degenerative disease^16,17^. Studies have suggested that maintenance of both Sirtuin 1 and Sirtuin 6 proteins are protective in vascular calcification^18^. Therefore, development or finding Sirtuin 1 and Sirtuin 6 regulating reagents may provide new strategies to prevent and improve vascular calcification.

Nicotinic acid (Niacin) is one of the lipid-lowering regents used in statin-intolerant patients as an alternative and is shown to reduce mortality in AS patients^19^. Previous studies indicated that niacin may be protective for cardiovascular diseases such as reducing monocyte/macrophage inflammatory responses, regulating oxidative stress and alleviating hyperlipidemia^20-22^. In vivo, niacin is converted to nicotinamide adenine dinucleotide (NAD^+^), which is reported to play an important role in VSMC fate, is required for the activity of numerous multifunctional enzymes, including Sirtuin 1 and Sirtuin 6^23,24^. Recently, niacin supplement is reported to attenuate pulmonary artery hypertension^25^. In addition, niacin supplement can regulate VSMC phenotype and protect against abdominal aortic aneurysm via increasing NADP-dependent SIRT1 activity in VSMC^26,27^. However, it is not known whether niacin can abrogate vascular calcification.

Herein, we evaluated if niacin treatment could counteract arterial calcification in both CKD and VD3-induced vascular calcification model. Besides, we explored the potential mechanism and our results indicated that niacin may mitigate vascular calcification by upregulating Sirt1 and Sirt6 expression and activating VSMC autophagy. These findings provide new insights in the prevention and treatment of vascular calcification.

## Material and Methods

### Association between niacin and AAC based on NHANES analysis

National Health and Nutrition Examination Survey (NHANES) is a large-scale, nationwide cross-sectional survey aimed at assessing the nutritional and health status of the US population. The NHANES study uses a complex weighted sampling method to ensure the sample is representative. In this study, all analyses were conducted according to NHANES’s official data analysis guidelines. As AAC-related data is only available in the 2013-2014 survey cycle, we included only participants from the 2013-2014 cycle. Participants without niacin intake and AAC index data were excluded, resulting in a final sample size of 2897 participants.

Detailed baseline data for all participants were collected, including age, sex, race, education level, and annual household income, obtained directly from the statistical questionnaires. Experienced measurers conducted detailed measurements of each participant’s height and weight, and BMI was calculated to assess obesity status. Smoking and drinking statuses were self-reported by the participants. For hypertension diagnosis, experienced measurers took blood pressure readings for each participant, averaging the results of five measurements. An average systolic pressure greater than 140 mmHg and a diastolic pressure greater than 90 mmHg were considered indicative of hypertension. Additionally, participants reporting a history of hypertension or taking hypertension medications were also considered to have hypertension. All hematological data were obtained after participants had fasted for at least eight hours. Detailed measurements of participants’ lipid profiles and blood calcium and phosphorus levels were performed. To explore the relationship between niacin and abdominal aortic calcification (AAC), we used the RCS curve method, adjusting for confounding factors, to fit a curve illustrating the relationship between niacin intake and the risk of AAC.

### Animal studies

All animal experiments were approved by the Institutional Animal Care Committee at Nanjing First Hospital, Nanjing Medical University and complied with the ARRIVE guidelines. Ten-week-old male C57BL/6J mice were purchased from GemPharmatech Co., Ltd (Nanjing, Jiangsu province, China). Mice were housed in a customized pathogen-free room with an ambient temperature of 25℃ and a humidity between 30% and 70% and were exposed to 12-h light–dark cycles and fed with rodent food and adequate water. All animals used were healthy and immune-normal. Adenine diet-induced (Xie Tong Sheng Wu, China) mice CKD model was established as previously described^28^. VD_3_ overload-induced CV was performed by s.c injection of 100 µL VD3 (5.5×105 U/kg) (MCE, Shanghai, China) once a day in 16-week-old mice for three times as described^29^. In some experiments, CKD mice were treated with EX527 (5mg/kg/d, MCE, Shanghai, China) and OSS_128167 (10mg/kg/d, MCE, Shanghai, China) by intraperitoneal injection for 4 weeks^30,31^. EX527 was dissolved in 1% DMSO (in physiological saline) and then added to culture medium to reach a final concentration of 100 μM. OSS_128167 was dissolved in 1% DMSO (in physiological saline) and then added to PBS to reach a final concentration of 200 μM. Niacin was given via gavage at a dose of 600 mg/kg once a day at the beginning of adenine treatment.

### Cell culture

Primary rat aortic smooth muscle cells were prepared using explanting method as previously described^32^. Briefly, 7w mice were intraperitoneally euthanized using sodium pentobarbital (150 mg/kg) and aortic arteries were separated. Aortic segments were then cut into small pieces and grown on the culture dish with SMCM (Sciencell, #1101, Carlsbad, CA, USA) added into it at 37°C in a humidified incubator. 3 days later, primary vascular smooth muscle cells were migrated from the aortic explants. Cells at passage 4 to 8 were used for further study. In some experiments, cells were treated with niacin at different concentrations (10, 25, 50 mM) in the presence of calcifying medium. Cells were treated with EX527 (10 μM) or OSS_128167 (20 μM) in some experiments.

### Small interfering RNA (siRNA) infection

Primary rat aortic smooth muscle cells were seeded in the 6-well plate. Cells were transfected with SIRT1 or SIRT6 siRNA (Gemagene, Shanghai, China) when it reached a density of 80%-90% by using Lipofectamine 3000Transfection Reagent (Thermo Fisher Scientific) according to the manufacturer’s instructions. Scramble siRNA was used as negative control. The efficient of siRNA was examined by immunoblotting.

### ALP activity assay

Alkaline phosphate (ALP) activity was measured by using alkaline phosphate assay kit (Beyotime, #P0321S, Shanghai, China). Cells were collected in ice-cold 0.1% Triton X-100 PBS. Protein concentration was measured using a BCA Protein Kit (Yeasen, #20201ES76, Shanghai, China). Protein samples were mixed with p-nitrophenylphosphate (p-NPP) substrate then incubated in 37℃ for 10 min. The reaction was terminated with 3 M NaOH. ALP activity was measured at 405 nm and was calculated as unit/mg protein.

### Alizarin S red staining

For cell staining, vascular smooth muscle cells were seeded in 6-well plate. After used medium and reagents were removed, cells were fixed using 4% paraformaldehyde (PFA) for 15 min at room temperature. After that, 1% alizarin red solution (Solabio, PH=4.2, #G1452, Beijing, China) were added for 10 min at room temperature. After disposing of the solutions, cells were rinsed with ddH_2_O to eliminate excess dye and images were taken under an inverted microscope.

For the whole mount of aorta staining, aortic arteries were separated then fixed in water-free ethanol for 24h. after that, arteries were stained with 0.003% alizarin red solution (Solabio, PH=8.3, #G1450, Beijing, China) in 1% potassium hydroxide overnight. The aortic arteries were then washed with ddH_2_O at room temperature for 10 min three times. Aortic arteries were then rinsed in 2% potassium hydroxide and photographed by using an inverted microscope.

### Calcium content assay

Calcium content assay was performed using a calcium content assay kit (Leagene, #TC1023, Beijing, China) in accordance with the manufacturer’s instructions. Briefly, cell or tissues were homogenized and the supernatant was isolated via centrifugation. 200 microliter of Methyl thymol blue (MTB) solution was mixed with 2.5μl samples and then incubated at room temperature for 10 min. the absorbance was examined at 610 nm by using a microplate reader. BCA Protein Assay (Yeasen, #20201ES76, Shanghai, China) was performed to assess total protein concentration. The relative calcium content normalized to the protein concentration was marked as μg/mg protein.

### Prediction of niacin targeted proteins

The SMILES formula of niacin was retrieved in the PubChem database (https://pubchem.ncbi.nlm.nih.gov/) and imported into the SwissTargetPrediction tool (http://swisstargetprediction.ch/) to predict the potential binding targets of niacin, and a total of 7 targets were obtained, and there was a possibility of binding.

### Molecular docking

The molecule structure of niacin was obtained from the PubChem database, and the structure of SIRT family proteins were obtained from the AlphFold database (https://alphafold.ebi.ac.uk/). Then, the active pockets of these 7 proteins were predicted by the CavityPlus tool (http://pkumdl.cn:8000/cavityplus/index.php), and finally the niacin and these active pockets were docked by AutoDock Vina, and the one with the lowest binding energy was selected for visualization.

### Von-Kossa staining of aortic sections

Aortic section von-Kossa staining was performed using a Von-Kossa Stain Kit (abcam, #ab150687, Cambridge, UK) according to the manufacturer’s protocols. In brief, paraffin aortic sections were deparaffinized and hydrate in distilled water. Then, section slides were incubated in Silver Nitrate Solution (5%) for 30 min with exposure to ultraviolet light. Distilled water was used to rinse the slides for 5 min. After that, sections were incubated with Sodium Thiosulfate Solution (5%) for 3 min. Next, aortic sections were rinsed in running water for 2 min followed by 2 changes of distilled water followed by incubation in Nuclear Fast Red Solution for 5 min. Then, aortic sections were rinsed in running water for 2 min followed by 2 changes of distilled water and dehydrate very quickly in 3 changes of fresh absolute alcohol and sealed with neutral resin (Sinopharm Chemical Reagent Co., Ltd, #10004160, Shanghai, China).

### Western blotting analysis

Cell or tissue were lysed with RIPA buffer containing proteinase (Yeasen, #20124ES03, Shanghai, China) and phosphatase inhibitors (Yeasen, #20109ES05, Shanghai, China). Then, protein concentration was determined using a BCA Protein Assay Kit (Yeasen, #20201ES76, Shanghai, China). 40 μg of protein was loaded in SDS-PAGE and then was transferred to PVDF membranes. Next, membranes were blocked in 5% defatted milk TBST solution followed by three times washing with TBST and incubation with primary antibodies diluted in Western Blot Primary Antibody Dilution Buffer (#WB500D, NCM Biotech, Suzhou, China) overnight at 4℃. The next day, membranes were washed with TBST followed by incubation with the accordant HRP-conjugated secondary antibody. The blots were visualized by the ECL image system and qualified using the image J software.

All the primary and secondary antibodies used in western blotting are listed as follows: anti-Runx2 (1:1000, #12556S, CST, MA), anti-Osteopontin (1:1000, #ab63856, Abcam, Cambridge, UK), anti-SIRT1 (1:1000, #13161-1-AP, Proteintech, Wuhan, China), anti-SIRT6 (1:1000, #67510-1-lg, Proteintech, Wuhan, China), anti-β-Actin (1:100000, #AC026, Abcolonal, Shanghai, China), HRP-conjugated Goat anti-Rabbit IgG (H+L) (1:10000, #ZB-2301, ZSGB-BIO, Beijing, China), HRP-conjugated Goat anti-Mouse IgG (H+L) (1:10000, #ZB-2305, ZSGB-BIO, Beijing, China).

### Statistical analysis

All results are presented as mean **±** SEM and all statistical analysis was performed by using GraphPad Software (GraphPad Software, Inc, USA). Differences between two groups were analyzed by Student’s *t* test. Multiple group datasets were analyzed by one-way ANOVA followed by followed by Tukey post-hoc tests. Biological experimental replicates between each group were shown in figure legends, and *p <* 0.05 was considered significant.

## Results

### 1. Niacin intake is negatively correlated with abdominal artery calcification

In this study, we included 2897 participants, of which 897 had abdominal artery calcification (AAC) and 2030 did not. As shown in Supplementary Table S1, baseline data indicated that participants in the AAC group were older than those in the non-AAC group. There was no significant difference in gender composition between the AAC and non-AAC groups. We found that the proportion of participants with an annual household income of less than $2000 was significantly higher in the AAC group compared to the non-AAC group. Additionally, the proportion of obese and overweight participants was significantly higher in the AAC group. Notably, the proportion of smokers was significantly higher in the non-AAC group, possibly due to the younger age of participants in this group. However, there was no significant difference in the proportion of drinkers between the two groups. Given that AAC and systemic arterial stiffness and calcification can lead to elevated blood pressure, the proportion of participants with hypertension was significantly higher in the AAC group. Since AAC can be partly due to abnormal calcium and phosphorus metabolism, we found that eGFR was higher in the AAC group compared to the non-AAC group. Furthermore, there were significant differences in lipid profiles between the two groups, as detailed in Supplementary Table S1. After adjusting for age, sex, race, household income, education level, obesity, smoking, drinking, hypertension, eGFR, blood calcium, and phosphorus levels, we further explored the relationship between dietary niacin intake and the risk of AAC. Using the RCS curve method, we performed a linear fit of this relationship. Our results suggested that as niacin intake increased, the risk of AAC significantly decreased, showing a linear negative correlation between the two (Fig. 1A).

**Figure 1.**
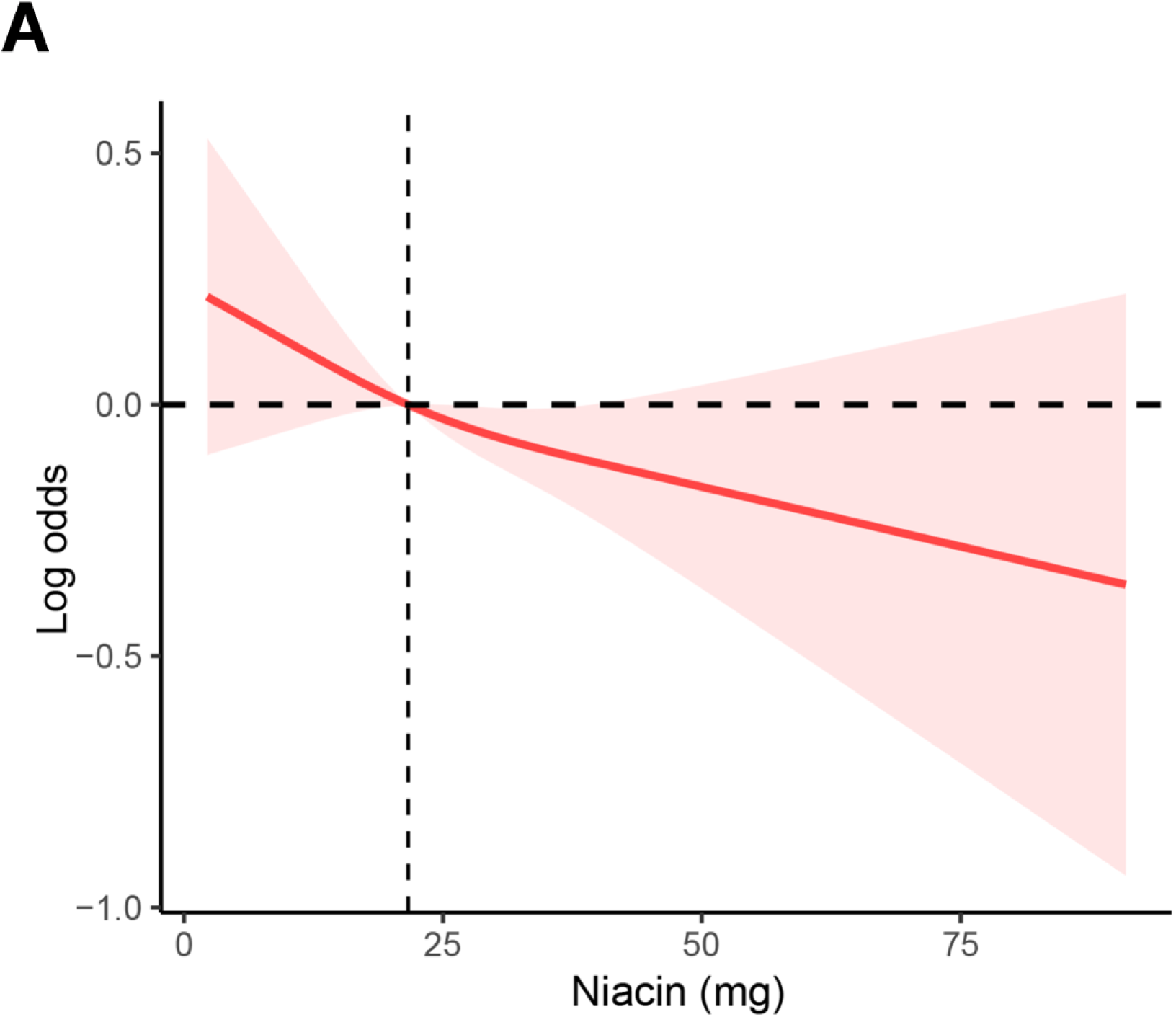
RCS analysis of the association between niacin intake and AAC.

### 2. Niacin attenuates calcification and senescence of vascular smooth muscle cell

To explore if niacin treatment affects the progression of vascular calcification. Primary rat aortic smooth muscle cells (RASMCs) were cultured in calcified medium (CM) containing high phosphate (2.6 mmol/L) to induce calcification. 0.1, 0.2 or 0.5 mmol/L niacin was added in the calcified medium^28^. Of note, CM-cultured VSMC exhibited obvious osteogenic phenotype as evidenced by significant upregulation of osteogenic markers Runx2 (runt-related transcription factor 2) and osteopontin (OPN) (Fig. 2A) and marked increasement of Alizarin red S staining (Fig. 2B). Intriguingly, niacin treatment significantly alleviated CM-induced VSMC calcification at day 14 as confirmed by qualification analysis of Alizarin red S staining (Fig. 2B). We also found that the expression of osteogenic markers including Runx2 and OPN were remarkably downregulated in VSMC treated with niacin compared with the untreated controls in a dose-dependent manner (Fig. 2A). Consistently, calcium content assay and alkaline phosphatase (ALP) activity assay showed that niacin treatment reduced both calcium content (Fig. 2C) and ALP activity (Fig. 2D).

**Figure 2.**
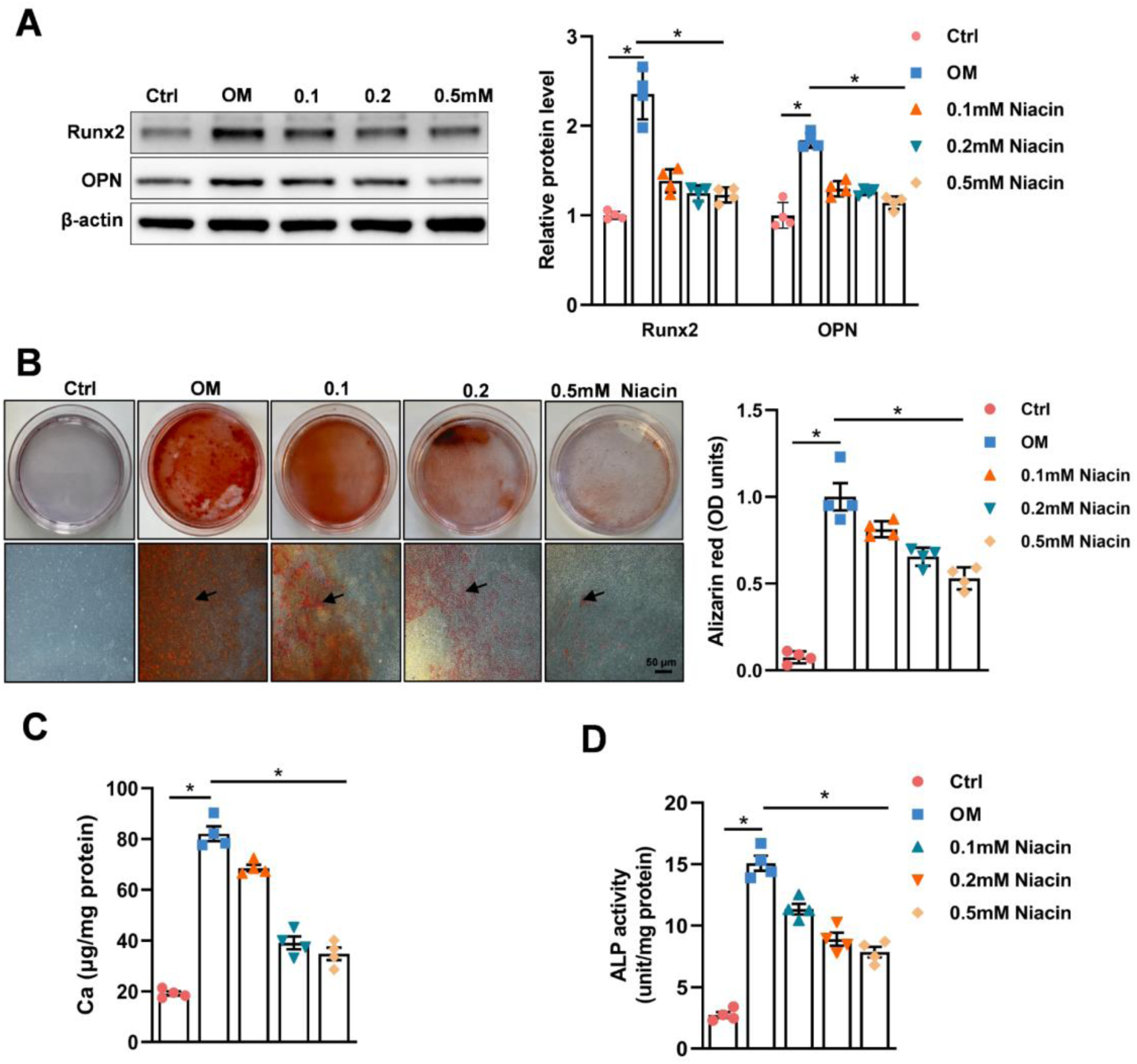
Niacin inhibits rat aortic vascular smooth muscle cell calcification. Rat vascular smooth muscle cells were incubated with growth medium (Ctrl), or calcifying medium (CM) supplemented with or without niacin with different concentration (0.1, 0.2 and 0.5mM) (n=4) for 3d or 14d. (A) Western blot was used to examine the expression levels of Runx2 and OPN after stimulation for 3 days. (B) Representative images showing cells stained with Alizarin red solution at day 14. Scale bar = 50μm. (C) Quantitative analysis of calcium content using a Ca assay kit. (D) quantitative analysis of ALP activity of RASMCs. * *p* < 0.05, ** *p* < 0.01. One-way ANOVA followed by Tukey’s post hoc test.

Since high phosphate stimulation is documented to induce VSMC aging, which is critical in the pathological process of VC^33,34^. We next examined the effect of niacin treatment on vascular smooth muscle cell senescence. Immunoblotting results indicated that niacin supplement significantly reduced cellular senescence-associated markers (p53 and p21), which parallels its previous impact on vascular calcification (Supplementary Fig. S1A). Moreover SA-β-gal staining (Supplementary Fig. S1B) and immunofluorescence staining (Supplementary Fig. S1C) revealed that osteogenic condition stimulated VSMC senescence was inhibited by niacin treatment. In addition, in vivo experiment showed aortic arteries from CKD mice expressed increased levels of cellular senescence-associated markers and niacin treatment greatly alleviated VSMC senescence via immunoblotting (Supplementary Fig. S1D) and immunofluorescence staining (Supplementary Fig. S1E). In all, these results demonstrate that niacin inhibits VSMC senescent phenotype switch and mineralization under osteogenic condition.

### 3. Niacin inhibits aortic calcification in C57BL/6J mice

Due to the fact that vascular calcification is usually found in CKD patients. We further assess the effect of niacin on CKD-induced vascular calcification. C57BL/6J mice were randomly assigned into chew diet, adenine diet and niacin group. Mice were fed with an adenine diet to induce CKD (Fig. 3A) and subsequent vascular calcification as previously documented^35^. CKD mice showed reduced body weight and increased serum levels of blood urea nitrogen, creatinine and phosphate compared with mice in chew diet group, yet no significant differences were observed between mice from adenine group and niacin group in serum levels of blood calcium, creatinine and phosphate before 12 weeks (Supplementary Table S2). Hematoxylin-eosin (H&E) staining of the kidney suggested identical renal injury between CKD mice and CKD mice supplemented with niacin (Supplementary Fig. S2A). Interestingly, we observed obvious difference in body weight and body urea nitrogen between adenine and niacin group in the middle and late stage of CKD (Supplementary Table S2). We next performed Micro-CT analysis to assess the extent of aortic calcification. Of interest, Micro-CT analysis indicated that niacin-supplemented mice exhibited a less calcification area compared with saline-treated mice (Fig. 3B). coordinately, Alizarin red S staining of aortas showed niacin supplement alleviated CKD-induced aortic calcification (Fig. 3C). As anticipated, von Kossa staining (Fig. 3D) and Alizarin red S staining (Fig. 3E) of aortic sections also showed niacin treatment reduced vascular calcification. Immunoblotting results demonstrated protein expression of calcifying markers were observably decreased in aortic arteries dissected from niacin group than that from adenine group (Fig. 3F). Moreover, we found niacin treatment significantly reduced calcium content in the aortic arteries (Fig. 3G). Finally, serum ALP activity was tested to show the inhibitory effect of niacin on VC (Fig. 3H). Collectively, these results show that niacin supplement neutralized CKD-induced VC in mice.

**Figure 3.**
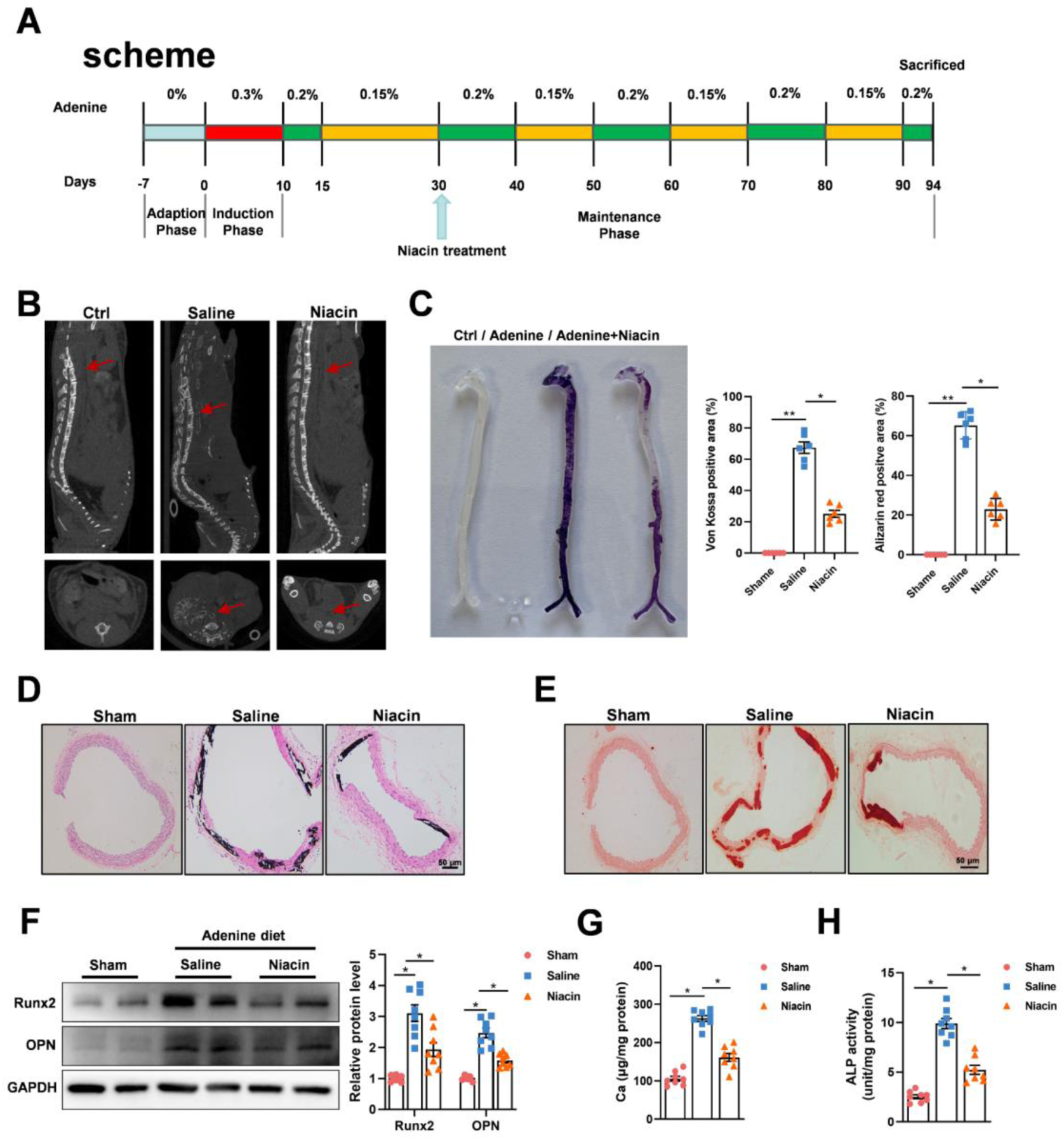
Niacin supplement alleviates CKD-induced vascular calcification in C57BL/6J mice. Adenine diet-induced CKD mice were treated with niacin (600mg/kg) via gavage for 12 weeks (n=8). (A) Scheme of the construction of the CKD-associated vascular calcification model and niacin supplement (B) Aortic calcification (red arrow) was examined by micro-CT. (C) Mineral deposition was detected by alizarin red staining of whole mount of aorta. Representative images showing aortic arteries staining with alizarin red. (D and E) Representative images of alizarin red and von Kossa staining of aortic sections. Scale bar = 50μm. (F) Representative western blot analysis and quantification of Runx2 and OPN protein levels in arteries from different groups. (G) Calcium content of aortas was measured. (H) Quantification of ALP activity. Data are shown as mean **±** SEM. \**p* < 0.05, \*\**p* < 0.01, N.S. not significant. One-way ANOVA, Tukey’s HSD post hoc test.

Since some clinical research indicated that niacin intake may exert beneficial effects on CKD^36^. To exclude that potential protective role of niacin. We used another vascular calcification model in C57BL/6J mice by injecting VD_3_ (Supplementary Fig. S4A) while treating them with saline or niacin for 6 weeks. We observed no significant alterations in body weight and serum mineral metabolites (Supplementary Table S4). Similarly, we observed less calcification area in niacin treated mice by Alizarin red S staining of whole aortas (Supplementary Fig. S4B). Of note, von Kossa staining (Supplementary Fig. S4C) and Alizarin red S staining (Supplementary Fig. S4D) of aortic sections also showed niacin treatment alleviated VD_3_ overload-induced vascular calcification. Moreover, immunoblotting result also indicated niacin treatment downregulated osteogenic markers in calcified aortic arteries (Supplementary Fig. S4E). Taken together, the above data suggested that niacin could mitigate medial arterial calcification.

### 4. SIRT1 and SIRT6 signal participate in VSMC calcification inhibition by niacin

To find out underlying targets of niacin in VSMC calcification. We first performed RNA-sequencing using arteries from adenine (VC) or niacin (VC + niacin) group (n=3 for each group). Whole transcriptome analysis identified 1834 upregulated and 1043 downregulated genes (fold change >2, P < 0.05) (Supplementary Fig. S4A). Kyoto Encyclopedia of Genes and Genomes (KEGG) analysis showed that downregulated genes in niacin group were enriched in pathways associated with bone mineralization and calcium-binding (Supplementary Fig. S4B). Gene set enrichment analysis (GSEA) validated the reduced expression of calcification (Supplementary Fig. S4C) and cellular senescence (Supplementary Fig. S4D) associated genes in niacin-treated vessels and upregulation of genes were enriched in nucleotide metabolism (Supplementary Fig. S4E) and longevity regulating (Supplementary Fig. S4F) pathway. Further, to find direct binding target of niacin, SwissTargetPrediction tool was used. Among 5 potential protein family candidates, the cellular longevity pathway (Supplementary Fig. S4F) highly correlated Sirtuin family was identified (Fig. 4A). Subsequently, we performed molecular docking to further investigate the interaction between niacin and 7 Sirtuin members. The docking results showed that SIRT6 and SIRT3, which are previously reported to be implicated in vascular calcification^37,38^, are most likely to directly bind to niacin (Fig. 4B). However, accumulating evidences have indicated that SIRT1 can also be regulated by niacin in cardiovascular pathologies^39,40^. Thus, we next examined if niacin regulated SIRT1, SIRT3 and SIRT6 under calcifying condition. Immunoblotting revealed that niacin treatment partly reversed OM-induced downregulation of SIRT1 and SIRT6 in VSMC without affecting the expression of SIRT3 (Fig. 4D, Supplementary Fig. S5A). To further determine the role of SIRT1 and SIRT6 signaling in niacin inhibited RASMCs calcification. We incubated RASMCs with SIRT1 inhibitor (EX527) and SIRT6 inhibitor (OSS_128167). Of interest, immunoblotting result showed that EX527 or OSS_128167 treatment diminished niacin-induced downregulation of Runx2 and OPN in smooth muscle cells (Fig. 4E). In addition, Alizarin red S staining indicated that either EX527 or OSS_128167 incubation partly blocked niacin inhibited VSMC calcification (Fig. 4F). Moreover, calcium content assay and alkaline phosphatase (ALP) activity assay also showed that EX527 or OSS_128167 treatment reversed niacin reduced calcium content (Fig. 4G) and ALP activity (Fig. 4H).

**Figure 4.**
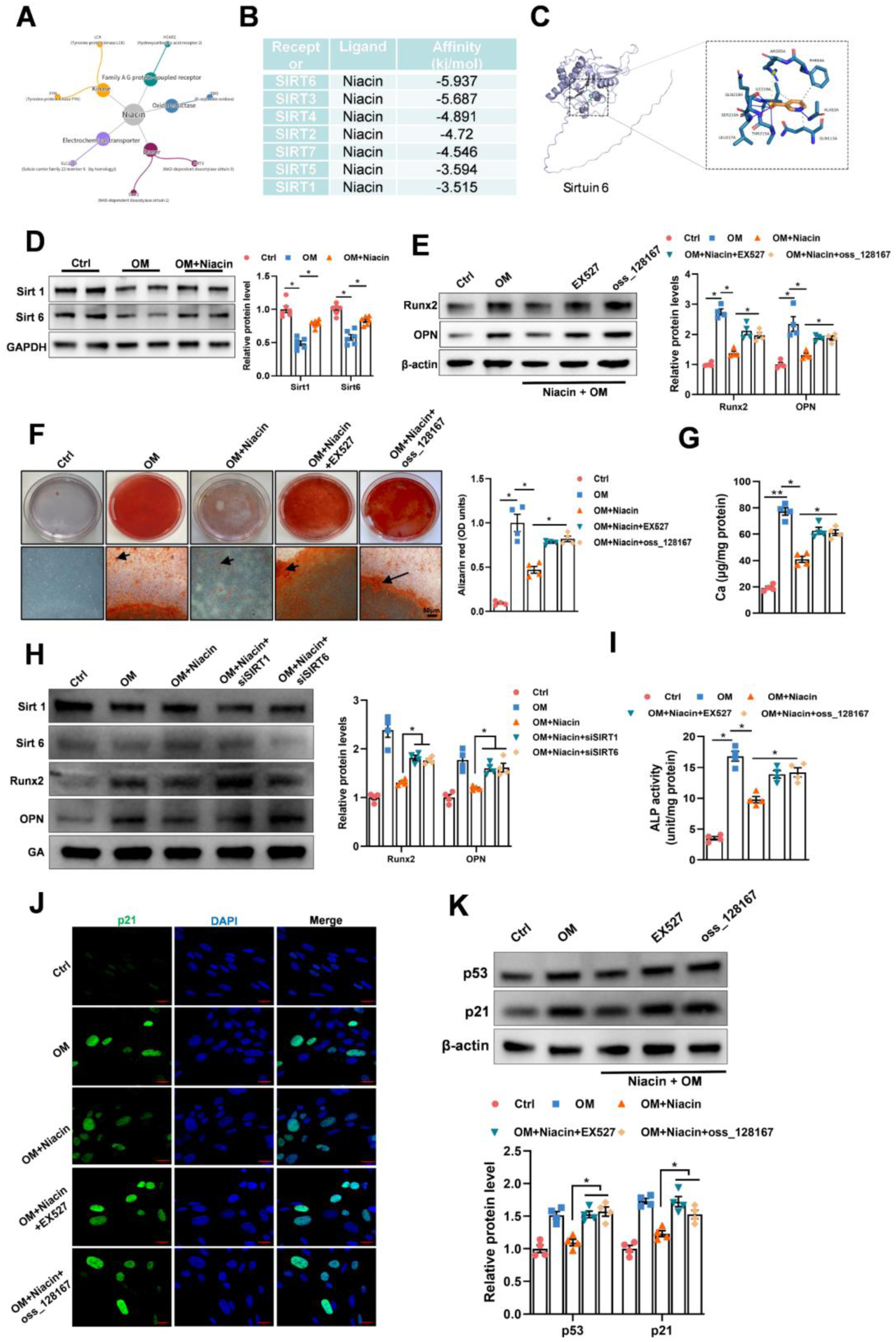
Niacin inhibited vascular calcification associated with SIRT1 and SIRT6 signaling. Pharmacological inhibition of SIRT1 or SIRT6 counteracted the beneficial effect of niacin on vascular smooth muscle cell calcification. Rat vascular smooth muscle cells were incubated with growth medium or calcifying medium in the presence of niacin or not (n=4). (D) Representative images and quantification of western blots for the expression levels of SIRT1 and SIRT6. RASMCs were treated with SIRT1 inhibitor EX527 or SIRT6 inhibitor OSS_128167 in the presence of CM and niacin. (E) Representative western blots for the protein expression of Runx2 and OPN. (F) Representative images showing cells stained with Alizarin red solution at day 14. Scare bar = 50μm. (G and H) Quantitative analysis of calcium content or ALP activity. (I) RASMCs were transfected with SIRT1 and SIRT6 siRNA in the presence of niacin and OM, representative images of western blots for the expression of Runx2 and OPN are shown. (J) Representative image of immunofluorescence staining in RASMCs. Scare bar = 20μm. (K) Representative immunoblotting image and quantitative analysis for the expression of p53 and p21 in RASMCs. Data are shown as mean **±** SEM. \**p* < 0.05, \*\**p* < 0.01, N.S. not significant. One-way ANOVA, Tukey’s HSD post hoc test.

To validated the effect of niacin on SIRT1 and SIRT6 signaling for its inhibitory role in vascular calcification. We performed RNA interference to knockdown SIRT1 and SIRT6 in smooth muscle cells in the presence of niacin. First, we examined the efficiency of RNA interference of SIRT1 and SIRT6 by western blotting (Supplementary Fig. S6A and S6B). Alizarin red S staining showed that SIRT1 or SIRT6 siRNA abrogated the protective effect of niacin on vascular smooth muscle cells. Besides, immunoblotting results suggested that niacin downregulated Runx2 and OPN expression levels were regained by SIRT1 or SIRT6 interference (Fig. 4I). Besides, Niacin improved VSMC senescence was counteracted after Sirt1 or Sirt6 inhibition by immunofluorescence (Fig. 4J) and immunoblotting (Fig. 4K). These data suggested that the SIRT1 and SIRT6 signaling are critical in niacin inhibited VSMC calcification.

### 5. Niacin attenuated aortic calcification in CKD mice via SIRT1 and SIRT6 signaling

To validate the requirement of SIRT1 and SIRT6 pathway in niacin-mediated vascular calcification inhibition in vivo. We supplemented CKD mice with EX527 or OSS_128167 or both together with niacin (Fig. 5A). First, we validated that niacin treatment reversed CKD-induced Sirt1 and Sirt6 downregulation in calcified vascular by immunoblotting (Fig. 5B). Micro-CT analysis revealed that EX527 or OSS_128167 supplement accelerated aortic calcification in niacin-treated CKD mice (Fig. 5C). Whole mount staining of aortas indicated SIRT1 and SIRT6 inhibition reversed niacin reduced vascular calcification (Fig. 5D). Accordingly, Von Kossa (Fig. 5E) and Alizarin red S (Fig. 5F) staining of aortic section also indicated niacin-alleviated vascular calcification through activation Sirt1 and Sirt6. Furthermore, we observed aortic expression of osteogenic-associated and senescence-associated markers were significantly upregulated after SIRT1 and SIRT6 inhibition (Fig. 5G). In all, these results demonstrated that the SIRT1 and SIRT6 signaling pathway are required in niacin attenuated vascular calcification.

**Figure 5.**
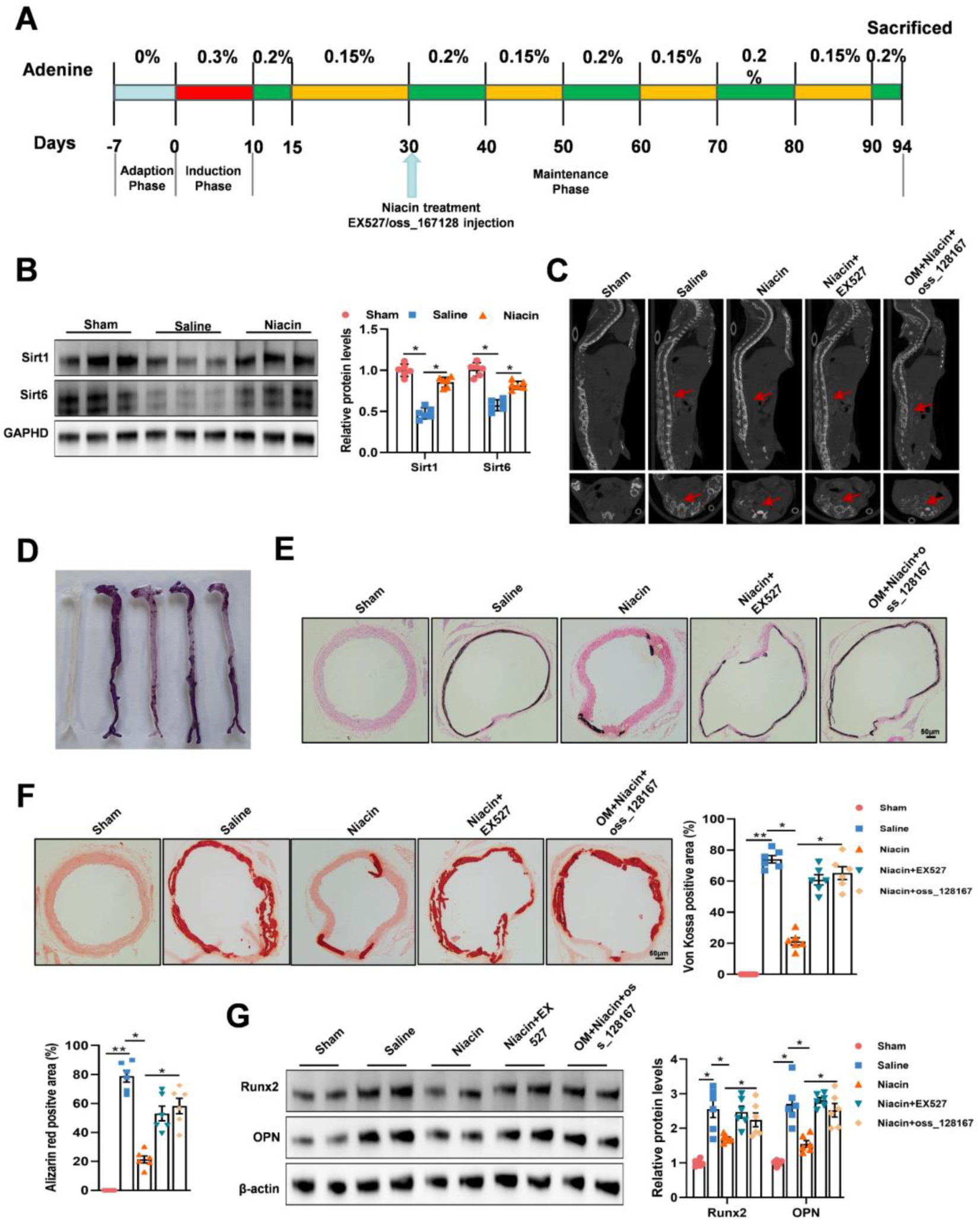
Effects of SIRT1 inhibitor and SIRT6 inhibitor on aortic calcification in mice with chronic kidney disease. Adenine diet-induced CKD mice were treated with niacin (600mg/kg) together with EX527 (5mg/kg/d, i.p.) or OSS_128167 (10mg/kg/d, i.p.) for 4 weeks. (A) Scheme of the construction of the CKD-associated vascular calcification model and EX527 or OSS_128167 supplement (B) Western blots for the expression levels of SIRT1 and SIRT6 of aortic arteries. (C) Representative images of aortic calcification (red arrows) are shown after examining by micro-CT. (D) Mineral deposition was detected by alizarin red staining of whole mount of aorta. Representative images showing aortic arteries staining with alizarin red. (E and F) Representative images of von Kossa and alizarin red staining of aortic sections. Scale bar = 50μm. (G) Representative western blot analysis and quantification of Runx2 and OPN protein levels in arteries from different groups. Data are shown as mean **±** SEM. \**p* < 0.05, \*\**p* < 0.01, N.S. not significant. One-way ANOVA, Tukey’s HSD post hoc test.

### 6. Activation of autophagy is required for the inhibition of vascular calcification by niacin

Previous studies have shown that autophagy plays important roles in vascular calcification^41^. Intriguingly, accumulating evidences have implicated that both SIRT1 and SIRT6 signaling may involve in the induction or regulation of autophagy^42,43^. Thus, we speculated that niacin inhibited vascular calcification may through SIRT1 and SIRT6-dependent autophagy pathway. To further test our hypothesis, we carried out western blot analysis to verify whether niacin treatment can influence autophagy in VSMC calcifying process. Interestingly, our results showed that OM treatment dramatically downregulated VSMC autophagy as indicated by the alteration of autophagy-associated markers such as Beclin1 and QSQTM1. This inhibitory effect of calcifying VSMC was partly reversed by niacin treatment (Fig. 6A). However, this niacin-induced activation of autophagy impact was diminished by SIRT1 or SIRT6 inhibitor supplement, suggesting niacin activated VSMC autophagy through the SIRT1 and SIRT6 (Fig. 6B). Rapamycin, a reagent that is known to enhance autophagy flux, was added in the culture medium of VSMC. As anticipated, rapamycin treatment evidently abrogated the calcifying effect of both EX527 and OSS_128167 (Fig. 6C and 6D). In vivo, we verified that niacin recovered CKD-reduced autophagy in vascular by immunoblotting (Fig. 6E). However, this effect was canceled after Sirt1 or Sirt6 inhibition (Fig. 6F and 6G). Apart from that, our immunofluorescence (Fig. 6H) and western blotting (Fig. 6I) results suggested that either SIRT1 or SIRT6 suppression acerbated niacin diminished VSMC senescence. All these results indicated that niacin treatment ameliorated vascular calcification via SIRT1 and SIRT6 mediated activation of VSMC autophagy (Fig. 7).

**Figure 6.**
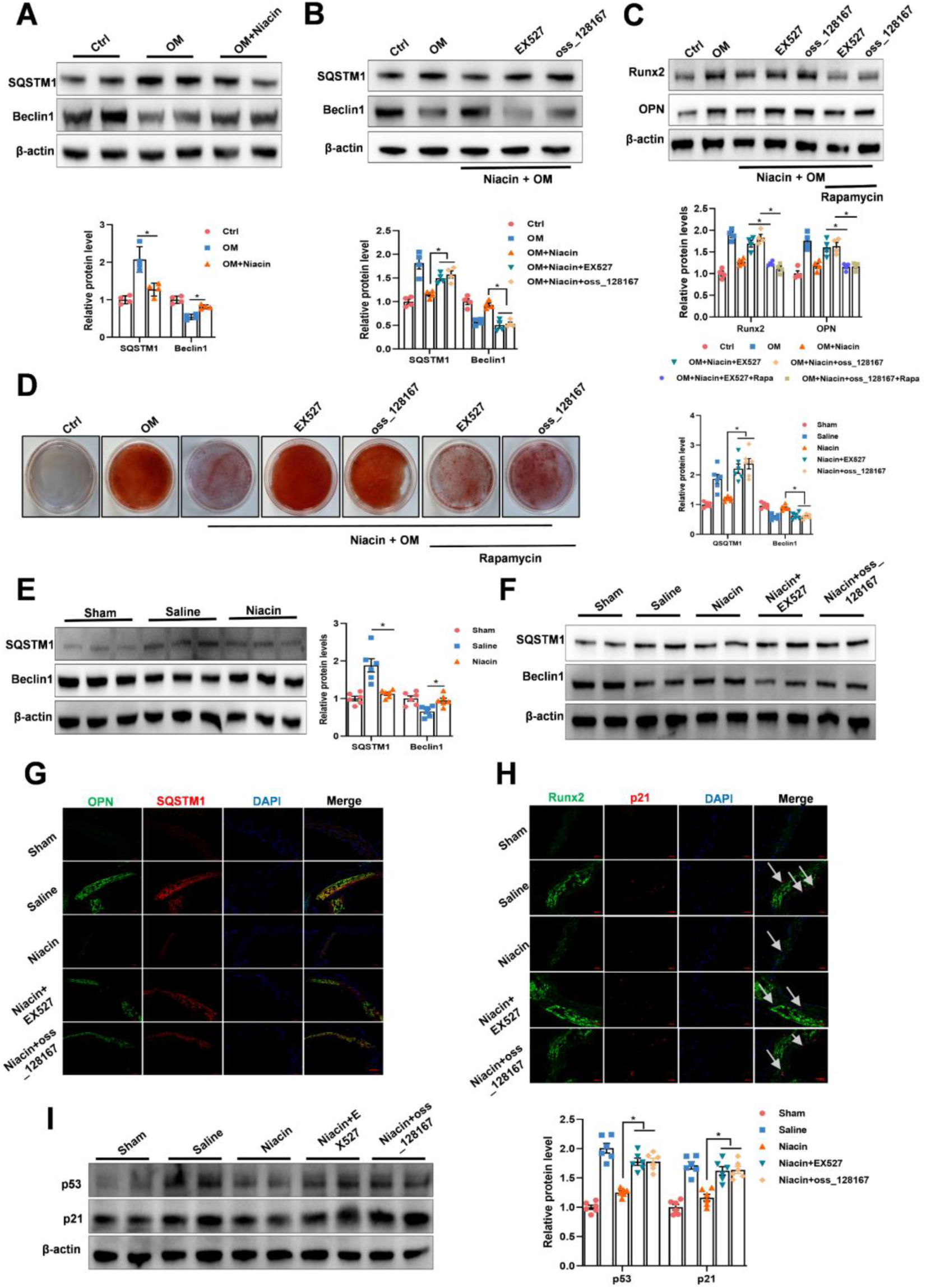
Niacin mediated inhibition of vascular calcification through SIRT1 and Srit6-mediated VSMC autophagy activation. (A) RASMCs were incubated with growth medium or calcifying medium in the presence of niacin or not. Autophagy-associated markers p62 and Beclin1 are examined by western blot. (B) Representative western blots for the expression levels of p62 and Belcin1 after EX527 and OSS_128167 treatment. (C) Representative western blots for the expression levels of p62 and Belcin1 after EX527 and OSS_128167 treatment together with rapamycin (200nM). (D) Representative images showing cells stained with Alizarin red solution at day 14. (E) and (F) Western blots for the expression level of vascular p62 and Beclin1. (G) Representative immunofluorescence images staining for p62 and Beclin1 of aortic arteries. Scare bar = 50μm. (H) Representative immunofluorescence images staining for Runx2 and p21 of aortic arteries. Scare bar = 20μm. (I) Representative image and quantitative analysis for the expression level of p53 and p21 of aortic arteries. Data are shown as mean **±** SEM. \**p* < 0.05, \*\**p* < 0.01, N.S. not significant. One-way ANOVA, Tukey’s HSD post hoc test.

**Figure 7.**
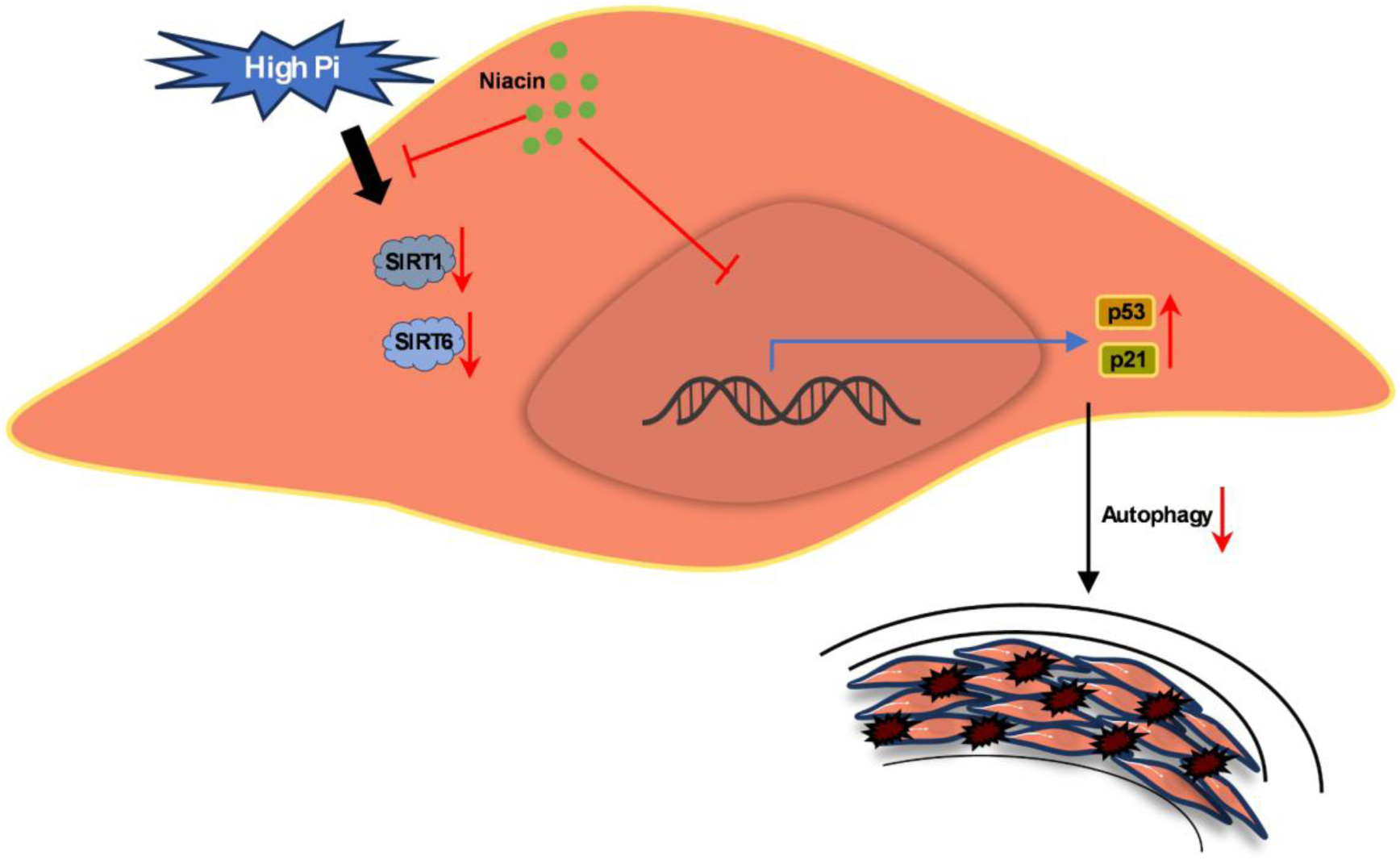
Niacin prevents vascular calcification by enhanced SIRT1 and SIRT6 signaling and promoted VSMC autophagy. SIRT1 and SIRT6 were downregulated in vascular smooth muscle cell (VSMC) under pathological conditions (e.g. CKD caused Hyperphosphatemia) promoting VSMC osteogenic transformation and vascular calcification. Niacin interacted with both SIRT1 and SIRT6 and maintained their levels. Apart from that, niacin treatment improved Sirtuin-mediated VSMC autophagy, thereby inhibiting vascular calcification.

## Discussion

Vascular calcification features ectopic mineral deposition of calcium and phosphate crystals in the tunica media of blood vessels that often displaying the characteristics of vascular aging and is considered as a complex and active process partially resembling bone mineralization^4,5^. Vascular calcification increases major adverse cardiovascular events in patients with chronic kidney diseases^44^. Although multiple mechanisms have already been recognized to contribute to the pathological process of vascular calcification, limited effective therapy is available. In the present study, we revealed that niacin treatment may be a promising inhibitor of vascular calcification. As an antihyperlipidemic agent, we demonstrated that niacin attenuated vascular calcification of vascular smooth muscle cell in a CKD and Vitamin D_3_-induced mouse model. To our knowledge, these data provide the first evidence that niacin treatment inhibited vascular calcification through modulating SIRT1 and SRIT6 and activating autophagy flux.

Arterial calcification is usually found in patients with chronic kidney disease and is correlated with aging in the vasculature system. Some common mechanisms are shared between aging and vascular calcification^45^. Intriguingly, previous studies have demonstrated that anti-aging reagents including resveratrol and irisin are protective in vascular calcification^46,47^. Niacin has been demonstrated to maintain endothelial nitric oxide to improve vascular aging^40^, and has emerged as an antioxidant that exhibits cardiovascular protecting effect^48,49^. Apart from that, niacin is reported to regulate inflammation and oxidative stress which are critical in the pathological progress of vascular calcification. Moreover, RCS analysis of the association between niacin intake and AAC indicated the protective role of high niacin diet in vascular calcification. Thus, we speculated that niacin could abrogate vascular calcification. Surprisingly, we found that niacin treatment inhibited the calcifying of rat aortic smooth muscle cells. Further, in vivo study indicated that niacin treatment obviously alleviated CKD and VD_3_ overload-induced vascular calcification. Additionally, a recent study has also implicated that niacin can protect against abdominal aortic aneurysm^26^. These data suggested that niacin may become a promising candidate for the treatment of vascular calcification in clinical applications since niacin exist in daily diet and has low toxicity yet strong and solid efficiency.

SIRT1 and SIRT6, two members of the Sirtuin family associated with cellular senescence, have been shown to inhibit vascular calcification. In calcified vascular, the expression of SIRT1 and SIRT6 are reduced^13,30^. Activation of both SIRT1 and SIRT6 may become a promising clue for treatment of vascular calcification. Niacin has been shown to activate SIRT1 in several studies^26,40,50^. Of note, we observed niacin treatment upregulated both SIRT1 and SIRT6 in calcifying VSMC. By using Sirt1 inhibitor EX527 and Sirt6 inhibitor OSS_128167, we found that EX527 or OSS_128167 treatment canceled niacin diminished vascular calcification in cultured VSMC and CKD mice. Our data indicated that both SIRT1 and SIRT6 are involved in the protective role of niacin reduced vascular calcification.

Accumulating evidences have suggested that autophagy, an important innate immune process that is responsible for the clearance of many pathogens and metabolic waste, plays a critical role in the regulation of phenotypic modulation of smooth muscle cell and vascular calcification. During the process of autophagy, QSQTM1 is downregulated and beclin1 is increased. Rapamycin, a classic autophagy activator, is reported to alleviate cell senescence, extend life span of animals and block vascular calcification. SIRT1 and SIRT6 are reported to be regulators of autophagy^51,52^. Notably, our results suggested that niacin significantly increased autophagy in vascular smooth muscle cells whereas EX527 and OSS_128167 obviously blocked it. Additionally, rapamycin treatment markedly prevented EX527 and OSS_128167 induced vascular calcification. Collectively, we first provide the finding that niacin treatment inhibits vascular calcification via promoting SIRT1 and SIRT6 mediated VSMC autophagy flux.

Although we verified the effective role of niacin in inhibiting vascular calcification. Limitations still exist in this study. As a reagent possesses multiple function including anti-inflammation, reducing oxidative stress and lowering hyperlipidemia. We only focused its impact on VSMC. Apart from that, although we illustrated that the effects of niacin on Sirt1 and Sirt6 and autophagy under calcifying state, the in-depth mechanism of how niacin regulates them awaits further exploration. Of note, a recent study has indicated the negative role of niacin metabolites in cardiovascular system, which may calls for more researches to carry on in the future^53^. In addition, even we performed two different vascular calcification models to exclude the impact of niacin on CKD. The current animal models are still not good enough to simulate vascular calcification in clinical patients. Finally, additional experiment is required to confirm the role of niacin on arterial calcification in SIRT1 and SIRT6 conditional knockout mice.

In conclusion, our study demonstrated that niacin acts as a novel reagent that inhibits vascular calcification. Furthermore, niacin decelerates CKD-induced vascular calcification via modulating SIRT1 and SIRT6 and affecting VSMC autophagy. Dietary proper niacin uptake may serve as a hopeful strategy for the treatment of vascular calcification. Our study has paves way for the prospective clinical trials to investigate the beneficial effect of niacin on arterial calcification of CKD patients.

## Acknowledgments

This work was supported by the National Natural Science Foundation of China (No.82000416, No.82170350) and Jiangsu Provincial Medical Innovation Center (CXZX202215).

## Conflict of interest

The authors declare that they have no known competing financial interests or personal relationships that could have appeared to influence the work reported in this paper.

## Author contributions

C.K. performed experiments, analyzed the data and drafted the manuscript; D.W. and Y.S. performed experiments and analyzed data. X.J., D.W., Y.S. and F.W. performed in vivo experiments. W.Z., J.L. and Y.G. analyzed the data and polished this manuscript. S.C. and Y.C. designed the experiment, revised the manuscript and supervised the study.

## Notes

### Competing Interest Statement

The authors have declared no competing interest.

